# Specifying cellular context of transcription factor regulons for exploring context-specific gene regulation programs

**DOI:** 10.1101/2023.12.31.573765

**Authors:** Mariia Minaeva, Júlia Domingo, Philipp Rentzsch, Tuuli Lappalainen

## Abstract

Understanding the role of transcription and transcription factors in cellular identity and disease, such as cancer and autoimmunity, is essential. However, comprehensive data resources for cell line-specific transcription factor-to-target gene annotations are currently limited. To address this, we developed a straightforward method to define regulons that capture the cell-specific aspects of TF binding and transcript expression levels. By integrating cellular transcriptome and transcription factor binding data, we generated regulons for four common cell lines comprising both proximal and distal cell line-specific regulatory events. Through systematic benchmarking involving transcription factor knockout experiments, we demonstrated performance on par with state-of-the-art methods, with our method being easily applicable to other cell types of interest. We present case studies using three cancer single-cell datasets to showcase the utility of these cell-type-specific regulons in exploring transcriptional dysregulation. In summary, this study provides a valuable tool and a resource for systematically exploring cell line-specific transcriptional regulations, emphasizing the utility of network analysis in deciphering disease mechanisms.

## Introduction

Transcriptional regulation significantly influences cellular function, development, responses to environmental factors, and pathologies. It plays a pivotal role in determining various phenotypes, including cancer. The human genome currently comprises over 1600 annotated transcription factors (TFs) that exhibit tight regulation and cell-type specificity (1). This highlights the importance of investigating regulation within specific cellular contexts. Extensive efforts have been directed towards understanding transcriptional regulation, resulting in the development of methods to construct regulons, representing sets of TF-target gene interactions that can be either direct or indirect. These methods vary in data sources, curation levels, and underlying hypotheses, lacking a unified annotation strategy.

One common method for creating regulons is manual literature curation. Several such databases like TRRUST (2), SIGNOR (3), and PAZAR (4) currently encompass regulons for approximately 800 human TFs. While this provides highly confident annotations, manual curation is hindered by resource-intensive collection and curation processes that are not easily scalable. To address this, text-mining techniques have been employed (5). These methods involve assigning confidence scores to sentences in abstracts of scientific articles. Highly confident sentences can then either be directly used to construct regulons or aid in the more rapid manual curation process. The outcome is the CollecTri resource, housing regulons for 1183 TFs and offering insights into regulatory interactions’ modes, such as activation or repression of target genes (6) making it valuable for estimating TF activity from downstream gene expression. However, while the CollecTri strongly benefits from its scale and high confidence of included curated sources, it lacks cellular context annotations.

Data-driven methods complement literature-driven approaches by integrating cellular context into Gene Regulatory Networks (GRNs) through two main method types: co-expression-based and TF-binding-based methods. Co-expression methods analyze transcriptomic data to identify genes interacting with a specific TF based on shared expression patterns (7–9), but they suffer from high false discovery rates due to non-causal correlations in gene expression (10, 11). In contrast, TF-binding methods utilize high-throughput chromatin immunoprecipitation followed by DNA sequencing (ChIP-Seq) data, identifying TF-DNA binding events in gene promoter and enhancer regions, with databases like ChIP-Atlas (12), and GTRD (13) providing such data. However, these databases often lack consideration of cellular transcriptional profiles when constructing regulons. For instance, ChIP-Atlas generates regulons for specific cell lines but it does not account for distinct gene expression patterns, potentially resulting in TF-target gene associations involving unexpressed genes within the studied cell line.

Owing to their partial overlap and complementary nature, various approaches aim to integrate data-driven and manually curated databases to offer regulons at multiple confidence levels. Databases like DoRothEA (14), RegNetwork (15), and CHEA3 (16), combine ChIP-Seq and co-expression-derived networks (ChEA3 and DoRothEA), alongside motif-based predictions and literature-curated sources (DoRothEA). RegNetwork contains predicted, literature-curated, and network-driven interactions, including protein-protein interactions, for both target genes and target microRNAs. Despite their comprehensive coverage, these resources still exhibit a noTable 3umber of false positive associations, primarily due to reliance on predictions, and they lack specific information about cell-type relevance.

In this context, we present a straightforward method to define regulons that capture the cell-specific aspects of both TF binding and target gene expression. Our approach uses data from ChIP-Seq and RNA-Seq experiments to construct regulons, and is easy to apply to any cell type with these data (17, 18). Here, we have applied it to four widely used cell lines with comprehensive ChIP-Seq data for a large number of TFs and functionally characterised them using various other types of biological networks. To validate our approach, we systematically benchmarked our regulons against existing resources (CollecTri, DoRothEA, ChIP-Atlas, TRRUST, RegNetwork) using the KnockTF database (19), revealing their comparable performance with the best methods. We also introduced filtering techniques that enhanced the predictive power of TF knockout benchmarking. Furthermore, through case studies, we showcased the ability of our regulons to identify relevant TF dysregulations in scRNA-Seq datasets from three cancer types, underscoring the significance of cell-type-specific transcription studies. Our tool and the regulons for K-562, Hep-G2, MCF-7, and GM-12878 are open access.

## Materials and Methods

### Data sources

Bulk RNA-Seq expression profiles of K-562 (ENCSR000CPH), GM-12878 (ENCSR843RJV), Hep-G2 (ENCSR019MXZ) and MCF-7 (ENCSR000CWQ) cells were obtained from ENCODE (17). Transcript expression values were averaged for isogenic replicates. Non-redundant ChIP-Seq data were acquired from the publicly available ReMap v.2022 database (18) and underwent filtering to exclude regions present in the ENCODE blacklist (20). Additional K-562-specific ChIP-Seq peak data for MYB were sourced from the GEO (GSE124541) database. The original peak calling pipeline (21) was employed using the hg38 reference genome. Further GFI1B data for K-562 cells were retrieved from the GEO (GSE117944) database (22) and lifted over to hg38. ATAC-Seq (ENCSR483RKN, ENCSR095QNB, ENCSR042AWH, ENCSR422SUG) and DNAse-Seq (ENCSR000EKS, ENCSR000EMT, ENCSR149XIL, ENCSR000EPH) data were collected from ENCODE as NarrowPeaks files.

### Mapping strategies

We introduced three distinct methodologies: “single TSS within 2 Mb” (S2Mb), “single TSS within 2 Kb” (S2Kb), and “multiple TSS within 2 Kb” (M2Kb), each characterized by different TSS selection criteria and varying window sizes around them (Table 1). The

**Table 1:** Overview of proposed approaches.

|  | S2Mb | M2Kb | S2Kb |
| --- | --- | --- | --- |
| Distance to TSS | [+1000; -1000] Kb | [+1; -1] Kb | [+1; -1] Kb |
| Selected TSS | Highest expressed isoform | Top 50% expressed isoforms | Highest expressed isoform |
| # transcripts per gene | Single | Multiple | Single |

TSS coordinates, as well as additional transcript and gene level annotations, were obtained using biomaRt v.2.48.3. for coding and non-coding Ensembl genes present in the bulk RNA-Seq profiles of the respective cell lines. Within these methodologies, we selected either the TSS coordinate of the highest expressed isoform (S2) or the TSS coordinates for the top 50% expressed isoforms (M2). It is important to note that we retained the most abundant TSSs (S2) or the top 50% most abundant TSSs (M2) for non-expressed genes to account for any potential modulation experiments.

To annotate TSSs of putative target genes with corresponding TF binding sites, we employed the bedtools v2.29.1 closest tool. Subsequently, distance filtering was applied as outlined in Table 1 to exclude unwanted interactions. This mapping process yielded a single association between a specific peak and its corresponding target gene for the S2Mb and S2Kb methods. In contrast, for the M2Kb approach, multiple associations were maintained between a peak and a gene due to the resolved isoform structure with a single peak per transcript but several transcripts per gene.

### Target gene annotation and enrichment analysis

We employed various binary features to characterise target genes, as detailed in Table 2. To annotate the promoters of target genes with ATAC-Seq and DNAse-Seq peaks within a 2000 bp distance threshold around the TSS, we used the GenomicRanges v.1.50.2 R package. For the motif annotation, we obtained PWMs from HOCOMOCO v.11 (23) and CIS-BP built 2.00 (24). To annotate TF motifs, we employed the “annotatePeaks” tool, and for enrichment analysis, we used the “findMotifsGenome” tool both within the HOMER v.4.11 software package. To analyse coexpression patterns between TFs and their target genes, we utilized WGCNA v.1.70-3 R package. We calculated coexpression measures for genes from single-cell RNA-Seq log-normalized count matrices obtained from K-562 cells with non-targeting sgRNAs, available at https://gwps.wi.mit.edu/. We established a correlation threshold of 0.1, resulting in the selection of the top 40% of correlated gene pairs.

**Table 2:** Overview of binary annotations of constructed regulons.

| Feature | Description |
| --- | --- |
| is_ppi | Whether there is a PPI between a gene and a given TF in STRING database (25) |
| is_coexpressed | Whether the co-expression of a gene with a TF is greater than 0.1 |
| network | Whether there is an interaction between a gene and a given TF in CRISPRi-derived networks from Morris et al. |

For network enrichment analysis, we employed separate logistic regression models for each binary characteristic (Table 2) with the glm function from the stats R package. This analysis considered presence in the regulon as the dependent variable, while gene expression and binary characteristics were included as covariates. As a random control, a permuted network was generated by shuffling target genes across TFs of the S2Mb regulon. The comparison of log2 odds ratio distributions was conducted using the Wilcoxon test with FDR correction.

### KnockTF Benchmarking

We utilized the benchmarking pipeline provided by the decoupler package at https://decoupler-py.readthedocs.io/en/latest/notebooks/benchmark.html. In essence, this pipeline first calculates transcription factor activities for TF knockout experiments using differential gene expression profiles from the KnockTF2 database (19); Fig. 2A). By leveraging prior knowledge about the perturbed TFs in knockout experiments, it then evaluates the predictive capacity of the networks using the area under the receiver operating characteristic (AUROC) and precision-recall curve metrics (AUPRC). Given that the true positive class is confined to the perturbations covered in the database, the metrics are computed using the Monte Carlo method. In each permutation, the negative and positive classes are balanced by randomly subsampling the former. For each specific cell line, we curated relevant knockout experiments from the KnockTF2 database, applying a filter to retain only high-quality experiments with a logFC of the perturbed TF greater than -1. GM-12878 was excluded from the analysis due to the lack of perturbational experiments. To ensure fairness in the comparison, we considered the intersection of sources (TFs) between the analyzed regulons and treated interactions in the CollecTri and DoRothEA regulons without prior knowledge of their mode of action (i.e., activator or repressor). The benchmarking pipeline was executed with default parameters.

**Figure 1:**
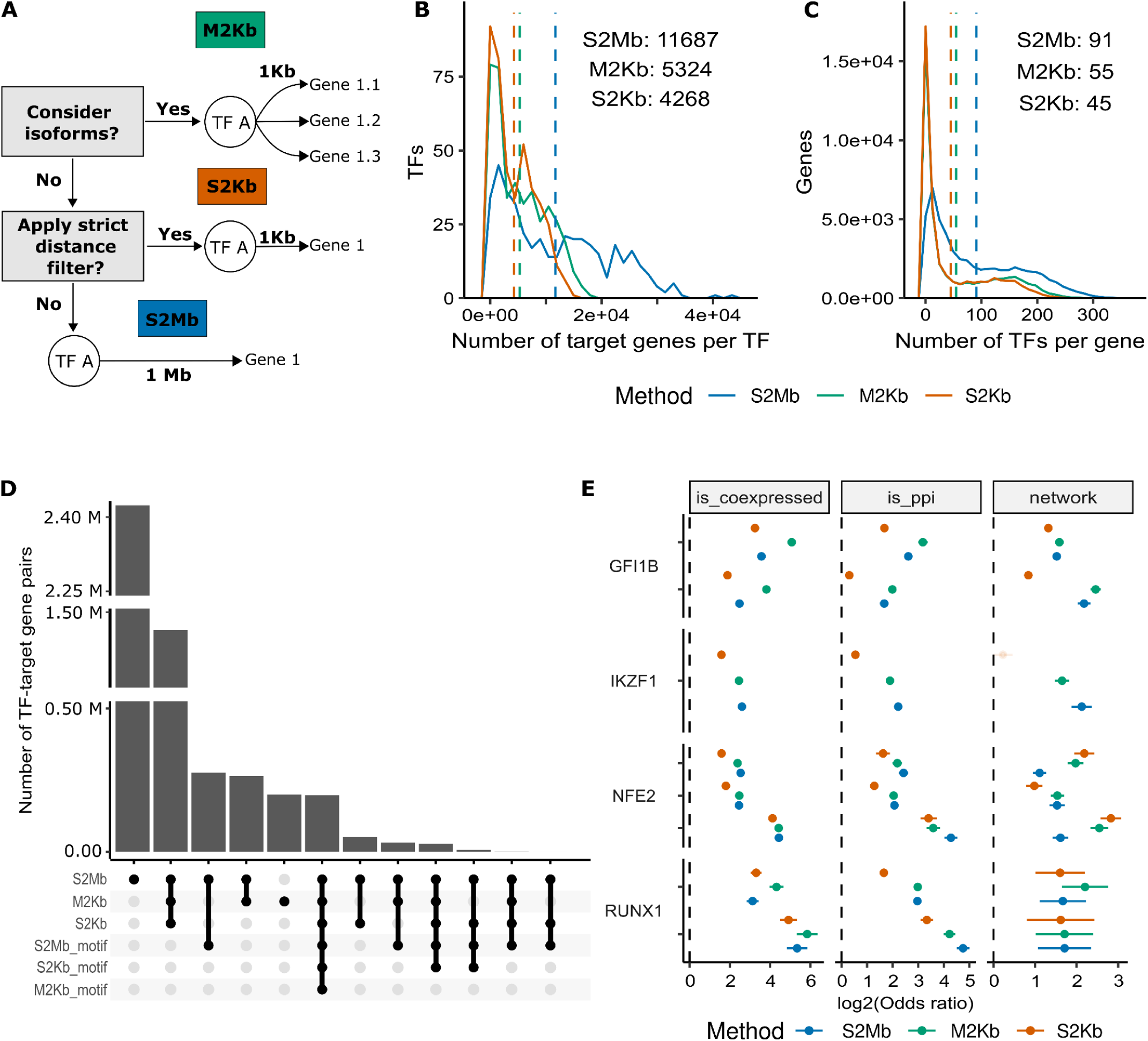
Overview of approaches and K562 regulons characteristics. (A) Schematic overview of proposed methods with specified distance cutoff and number of considered transcripts. (B) Distribution of a number of target genes per TF; dashed line shows per method median. (C) Distribution of a number of TFs per target gene; dashed line shows per method median. (D) Overlap of regulons with TF binding motifs; the top 12 overlapping groups are shown. (E) Enrichment of regulons for K-562 cell line in biological networks: coexpression networks from scRNA-Seq data (left), PPI networks (middle), trans-regulatory networks identified using STING-Seq technique (Morris et al. 2023) (right). Here, independent ChIP-Seq experiments result in multiple dots within a transcription factor.

**Figure 2:**
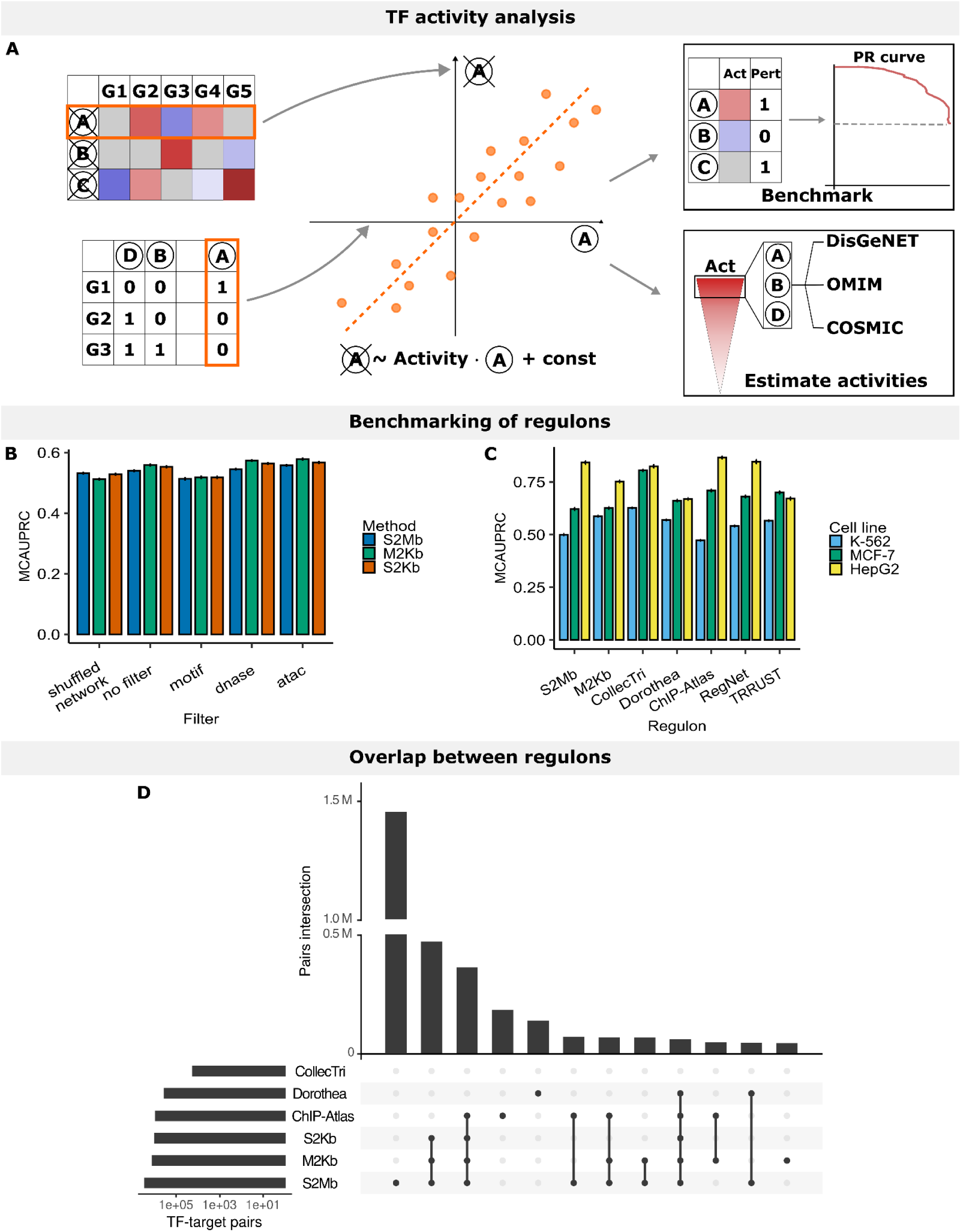
Overview of benchmarking procedure. (A) Schematic overview of the benchmarking procedure. Here, expression values from a TF knockout experiment and a regulon of interest (left) are used as inputs for activity estimation. Here different transcription factors are encoded using letters (e.g. A, B, D) and genes are referred to as Gx. We fit a univariate linear model to model gene expression as a function of TF regulations and estimate activities of transcription factors as regression coefficients of this model (middle). Then, in a benchmarking setting, we classify TFs into perturbed or non-perturbed and estimate its goodness (right upper) or, for the case studies, we perform activity-based ranking of TFs and an enrichment analysis (right bottom). (B) - (C) Average MCAUPRC metric represents the predictive power of a TF activity-based classifier built upon different regulons. (B) Comparison of filtering strategies for S2Mb, M2Kb, and S2Kb for K-562 regulons using knockout experiments from the KnockTF database. (C) Comparison of regulons in identifying TF perturbations in knockout experiments from KnockTF database across cell lines. (D) Overlap in the TF-target gene interactions between K-562 regulons from different resources; for ease of interpretation, the top 12 overlapping groups are shown.

### Single-cell RNA-Seq datasets

We acquired preprocessed single-cell data from breast cancer (26); GSE176078) and hepatoblastoma liver cancer (27); GSE180665) studies. For the hepatoblastoma dataset, we selected hepatocytes and a subset of neoplastic cells to ensure an equal number of cells in each group. Within the breast cancer dataset, we designated epithelial cells labelled as LumA_SC and Basal_SC as cancer cells, while no_scTYPER_call cells were identified as healthy controls, following the original study’s annotations. Differential expression analysis was performed using the Wilcoxon test within scanpy package. Genes with FDR below 0.01 were selected as differentially expressed genes (DEGs), without applying a log-fold change (logFC) cutoff. In the case of the AML dataset (28), we employed the results of the differential expression analysis provided by the authors.

### Activity estimations

TF activities in each case study dataset were estimated using the decoupler package’s univariate linear model (ulm) according to standard guidelines https://decoupler-py.readthedocs.io/en/latest/notebooks/dorothea.html. Briefly, these activities were computed as t-values through linear regression, where the expression estimates (specifically logFC values from DE analysis) were regressed against the “TF profile”. This profile is a signed adjacency matrix representing TF-target gene interactions, indicating activating interactions with 1 and repressive interactions with -1. Uncharacterized TF regulatory modes were treated as activators. A sign-aware network was applied for the CollecTri regulon due to its improved performance (6). Dysregulated TFs were assigned if their p-value was < 0.01 without applying a t-value threshold.

### Disease gene enrichment analysis and interpretation

The lists of activated TFs underwent enrichment analysis using the enrich tool from the gseapy package along with gene sets from DisGeNET (29), OMIM_Expanded (30) and KEGG_2021_Human (31). An additional set of COSMIC consensus genes (32) was obtained via a decoupler package.

## Results

### Collected data

We obtained ChIP-Seq data from the ReMap database (18) focusing on four cell lines: K-562, Hep-G2, MCF-7, and GM-12878 (Table 3). We associated transcription factors (TFs) with target genes in each dataset by identifying the gene closest to each ChIP-Seq peak, using two distance filters (2Kb and 2Mb) and two gene annotation strategies (single and multiple transcription start sites [TSS] per gene; see Methods; Fig. 1A). In the single TSS (S2) approaches, we linked the initiation site of the most highly expressed isoform of each gene with TF peaks identified via ChIP-Seq experiments. Conversely, the multiple TSS (M2) approach incorporated start sites from the top 50% expressed isoforms of each gene. Our strategy considered two distinct windows: one within ±1 Kb around the TSS, capturing TF binding within gene promoters and nearby enhancers, and a broader ±1 Mb window, enabling us to detect TF regulation from more distant enhancers. Hereafter, each TFs target gene set is called a regulon, while interactions refer to TF-target gene pairs.

**Table 3:**
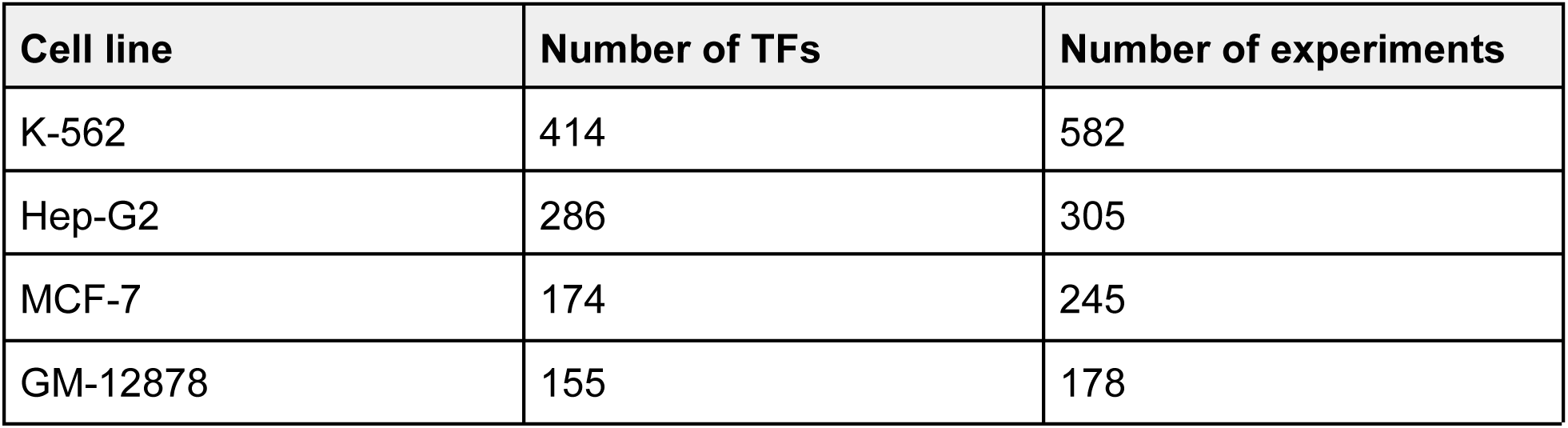
Summary statistics of ChIP-Seq data.

The S2Mb approach yielded double the number of target genes per TF compared to the M2Kb and S2Kb approaches (Fig. 1B-C, S1, Table 4). The M2Kb approach identified approximately 19% more target genes per TF than the S2Kb approach. Across cell lines, the median regulon size varied, ranging from 5, 190 target genes per TF in GM-12878 cells to nearly 8, 000 in Hep-G2 cells (Fig. 1B-C, S1, Table 4). Overall, the regulon size appeared to be higher in Hep-G2 cells.

**Table 4:** Median size of regulons, i.e. number of target genes per TF.

| Cell line | S2Mb | M2Kb | S2Kb |
| --- | --- | --- | --- |
| K-562 | 11687 | 5324 | 4268 |
| Hep-G2 | 15771 | 7997 | 6574 |
| MCF-7 | 14317 | 6081 | 5116 |
| GM-12878 | 11506 | 5190 | 4328 |

### Characterization of TF-target gene interactions

To characterize our regulons, we compared regulons and annotated promoters of target genes with TF binding sites (TFBS) from HOCOMOCO (23) and CIS-BP databases (24) using HOMER software (33). About 25% of TF-target gene interactions were exclusively identified by the S2Mb approach, which can capture TF binding outside promoter regions (Fig. 1D, S2). Approximately 10% of interactions displayed overlap with TF binding motifs in target gene promoters. Notably, S2Mb-identified interactions might potentially be depleted of motifs due to promoter-centric motif annotation strategy. We envision that the peak-centric motif annotation approach as well as improvements in TFBS annotation tools, along with the incorporation of a broader motif database, could potentially increase the number of confidently identified TF-target gene interactions.

Subsequently, we investigated the biological implications of the identified interactions by assessing their enrichment within various biological networks. Specifically, we examined co-expression networks, derived from single-cell RNA-Seq experiments (34), protein-protein interaction (PPI) networks sourced from the STRING database (25) and the trans-regulatory networks identified in CRISPRi experiments (35). Our results revealed a robust enrichment of our target genes in co-expression networks (Fig. 1E, S3A) and PPIs (Fig. 1E, S3B-E), often with higher values observed for the M2Kb approach (Fig. 1E, S3). Additionally, our regulons were enriched in experimentally derived trans-regulatory networks for four tested TFs (GFI1B, NFE2, IKZF1 and RUNX1) (with log2 odds ratios ranging from 1 to 3), thereby reproducing previously observed results (35). For the IKZF1 transcription factor, the S2Mb regulon exhibited a more pronounced enrichment within trans-networks compared to M2Kb, potentially due to its ability to capture distal enhancer-TF interactions.

### Benchmarking regulons using TF knockout experiments

Next, we evaluated the constructed regulons, starting by examining their overlap with comparable publicly available datasets: CollecTri, ChIP-Atlas and DoRothEA. Across most cell lines (K-562, MCF-7, and GM-12878), the highest degree of overlap was observed within our approaches (around 19% of interactions), followed by a significant overlap between our and ChIP-Atlas regulons (6%) (Fig. 2D, S4A, S4C). For Hep-G2 cells, the primary overlap occurred between our methods and the ChIP-Atlas database, accounting for 23.4% of interactions (Fig. S4B). Notably, DoRothEA and CollecTri regulons demonstrated limited agreement with other datasets (up to 5.5% of interactions). The CollecTri regulons exhibited limited overlap with others, primarily attributed to regulons’ size being two orders of magnitude smaller compared to other databases (Fig. 2D, S4). In summary, an average of 36.5% of interactions were shared across at least two datasets.

Considering the proportion of interactions unique for each regulon, we investigated their capacity to elucidate induced transcriptional alterations. We reasoned that regulatory TF-target gene interactions should manifest as changes in the expression of target genes following TF perturbation. Thus, we employed the decoupler GRN benchmarking pipeline and the KnockTF database (see Methods; Fig. 2A) to benchmark the methods. To address class imbalance between positive and negative classes, we employed a Monte-Carlo method to calculate classification metrics, i.e. area under precision-recall curve and area under receiver-operator curve (MCAUPRC and MCAUROC respectively). We compared the performance of our regulons with similar methods — TRRUST, DoRothEA, RegNet, CollecTri, and ChIP-Atlas — by benchmarking them separately for three cell lines (K-562, Hep-G2 and MCF-7).

In the absence of any filters, we observed a statistically significant difference (p <= 0.01, Wilcoxon test with FDR correction) in MCAUPRC across most cell lines and approaches, compared to the permuted network (Fig. 2B, S5A-B, S6A-C, S7A-C). On average, the S2Kb and especially the M2Kb approaches outperformed S2Mb, suggesting the usefulness of the stricter distance filter in mitigating false positive associations (Fig. 2B, S6C, S7C). We also explored filtering strategies based on expression (TPM > 0), open chromatin (DNAse-Seq, ATAC-Seq) and motif annotation to minimize spurious TF-target gene interactions. Remarkably, the inclusion of an open chromatin filtering step enhanced the regulons’ ability to predict the upstream TF perturbations, particularly benefiting the shorter-distance approaches (S2Kb and M2Kb) (overall p-value < 0.05, Wilcoxon two-sided test; Fig. S4-S6). In contrast, both the expression and motif filters contributed minimally to performance enhancement. Our results underscore the similar performance of the 2Kb methods, with the optimal outcomes achieved through the application of an open chromatin filter. Thus, we proceeded with M2Kb and S2Mb regulons, incorporating open chromatin filters, for subsequent comparisons with other datasets.

Following the same benchmarking procedure, we found that no single regulon consistently outperformed others in predicting TF perturbations. Performance metrics displayed notable variations within methods across different cell lines (Fig. 2C). CollecTRI outperformed other regulons in K-562 and MCF-7 cells, with average MCAUPRC of 0.63 and 0.80, respectively (Fig. S5C-D, S6D-E), attributed to its incorporation of interactions from text-mining and curated data sources. Interestingly, for Hep-G2 cells, ChIP-Atlas showed the highest score of 0.87 (Fig. S7D-E). The M2Kb approach consistently demonstrated solid performance across cell lines, yielding average MCAUPRC scores of 0.59, 0.63, and 0.75 for K-562, MCF-7, and Hep-G2, respectively (Fig. 2C, S5C-D, S6D-E, S7D-E). Notably, ChIP-Atlas outperformed our approaches in MCF-7 and Hep-G2 cells, despite employing similar mapping strategies, possibly due to the additional filtering of low-quality targets incorporated in our benchmarking pipeline (see Methods). In summary, while the M2Kb and S2Mb approaches did not rank among the top performers, they consistently demonstrated competitive performance on par with other data-driven regulons, with the key strengths lying in their transparency, user-friendliness, and efficient application compared to text-mining and curated resources.

### Regulon annotations identify the disrupted activity of TFs in disease

Transcription factors (TFs) can drive tumor growth and metastasis in a cancer-specific manner (36, 37). Thus, we hypothesised that regulons tailored to specific cell types could capture disease-specific transcriptional patterns (38). We utilized data from three single-cell RNA-Seq studies on distinct cancers: acute myeloid leukemia (AML) (39), breast cancer (26) and hepatoblastoma (27)). By analyzing specific cell types in both healthy and diseased cells, we estimated TF activities (40) employing four cell-line-specific regulons for K-562, Hep-G2, MCF-7, and GM-12878 derived using the M2Kb approach and ChIP-Atlas, as well as a generalized CollecTri regulon (Fig. 3A-D, S8A-D). Finally, we explored potential links between dysregulated TFs and cancer-associated genetic mutations through enrichment analysis using the databases DisGeNet (29), OMIM, and COSMIC (32) (Fig. 3E-G, S8E-G, S9, S10).

**Figure 3:**
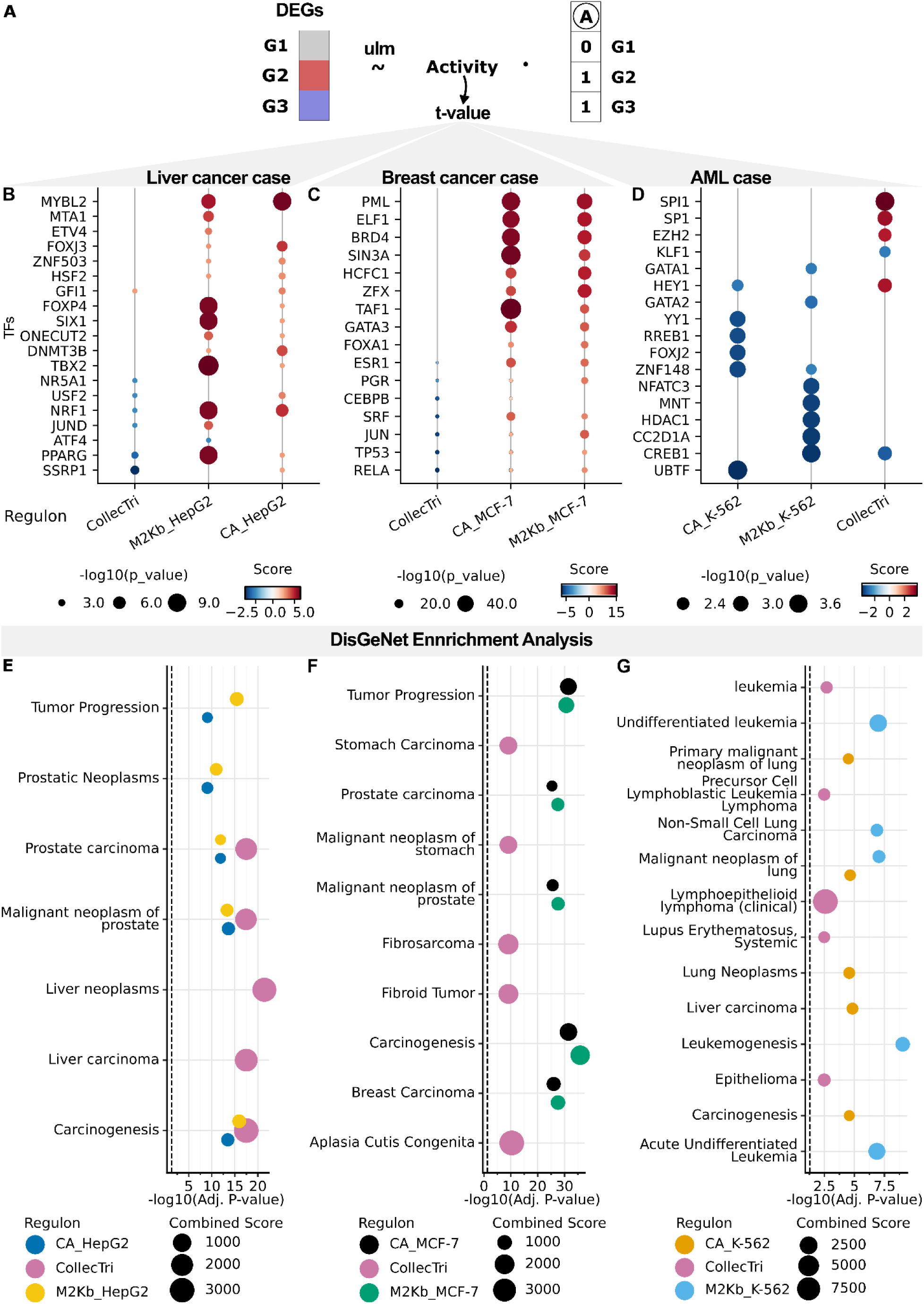
Case studies of detecting transcriptional dysregulation in cancers. (A) Schematic overview of activity estimation procedure. (B) - (D) Activity estimate for the top-ranked and manually selected TFs of each regulon for (B) hepatoblastoma (27), (C) lumenal type A breast cancer (26) and (D) leukemic progenitors (28). Here positive and negative activity scores mean activation and deactivation of TFs in malignant cells respectively. (E) - (G) Top 5 enriched terms of the enrichment analysis of dysregulated TFs in disease gene from DisGeNet database (29) for (E) hepatoblastoma, (F) lumenal type A breast cancer and (G) leukemic progenitor cells. Here “CA_cell line” refers to ChIP-Atlas regulons for the specified cell line and “M2Kb_cell line” refers to M2Kb regulons.

First, using gene expression levels to infer TF activity, we compared transcriptional profiles between neoplastic and healthy liver cells (27) and then performed gene set enrichments of dysregulated TFs (Fig. 3B, 3E). We observed that only CollecTri-detected dysregulated TFs were enriched in the gene sets associated with liver neoplasms in DisGeNet (Fig. 3E). In contrast, the dysregulated transcription factors identified by both the ChIP-Atlas and M2Kb-based Hep-G2 regulons were enriched in broader cancer-related terms in DisGeNet, such as “Cancerogenesis” and “Tumor Progression” (Fig. 3E). Among the activated TFs identified by the M2Kb regulon, six TFs (E2F1, ETV4, MTA1, MYBL2, NCOA3, and PPARG) have been used as prognostic markers for lower patient survival, and some other TFs were linked to liver cancer (41); Fig. 3B). Furthermore, we observed the activation of transcription factors associated with KEGG pathways such as “Transcriptional Misregulation in Cancer, “ “Hepatitis B” and “Non-alcoholic Fatty Liver Disease” (for both CollecTri and M2Kb) (Supplementary Table 1 - 2).

We then explored the potential to differentiate cancer types by analyzing transcriptional dysregulations in malignant epithelial cells from luminal A and basal types of breast cancer (26). In luminal A breast cancer, dysregulated TF sets detected by both M2Kb and ChIP-Atlas-based MCF-7 regulons were significantly enriched in breast and prostate cancer terms in DisGeNet (Fig. 3F, S9B, S10B). The CollecTri-based regulon identified a TF set enriched in fibrosarcoma and stomach carcinoma terms but not in breast cancer. Interestingly, cell type-specific activity estimates (M2Kb and ChIP-Atlas) showed heightened GATA3 activity in luminal A breast cancer, in contrast to reduced activity in basal type (Fig. 3C, S8B), aligning with its known role in promoting luminal activity and cell differentiation (42). Furthermore, established markers of luminal A breast cancer, such as FOXA1 (43), ESR1, and PGR, were activated in malignant cells of the luminal A cancer type (Fig. 3C). Conversely, all three regulon estimates indicated deactivation of these markers in basal-like cancer cells, consistent with the absence of ER, PR, and HER2 markers in triple-negative breast cancer (Fig. S8B).

Examining the transition from healthy hematopoietic stem cells (HSCs) to abnormal AML progenitor cells (39), we discovered that perturbed transcription factors in aberrant progenitors exhibited enrichment in leukemia and cancer-related signatures (Fig. 3G). Interestingly, TFs identified by the M2Kb-based GM-12878 regulon were enriched in lymphoma-associated but not leukemia-associated genes in DisGeNet, possibly due to the B-cell origin of GM-12878 (Fig. S8F). Within the M2Kb-based K-562 regulon context, deactivated TFs included GATA1, a switch factor for erythroid development (44, 45) and GATA2, crucial for hematopoietic development (46) and linked to myelodysplastic syndrome (47, 48); Fig. 3D). This suggests a potentially reduced commitment of analyzed progenitors to the erythroid lineage compared to HSCs, supported by CollecTri regulon-detected deactivation of the erythropoiesis driver gene KLF1 (Fig. 3D). Notably, SPI1, a known tumor suppressor that induces AML upon knockdown (49, 50), was activated in aberrant progenitors according to the CollecTri regulon (Fig. 3D).

Finally, we compared leukemic activated hematopoietic stem cells (HSCs) to their dormant counterparts categorized by mutational burden through CloneTrace (28). Using both M2Kb and ChIP-Atlas-based blood regulons (GM-12878 and K-562), we identified dysregulation in essential transcription factors involved in hematopoiesis and AML progression, such as MYB (51), MYC (52), SP1 (53), CREB1 (54) and YY1 (55) in activated leukemic HSCs (Fig. S8D). Intriguingly, CREB1 and YY1 were dysregulated according to the CollecTri regulon, however, marked as deactivated (Fig. S8D). While the transcription factor MYB was activated according to M2Kb-based GM-12878, it was not activated according to K-562 regulons, possibly due to the downregulated expression of MYB in K-562 cells (51). In general, transcription factors with perturbed activity from M2Kb-based GM-12878 regulon exhibited enrichment in leukemic and cancer-related terms, while other regulons detected TFs that were less specific to particular cancer types (Fig. S8G, S9F, S10F). Altogether, these analyses emphasize the utility of cell-type specific regulons for pinpointing transcription factor dysregulations in specific conditions consistent with previous observations (38).

## Discussion

Transcription is pivotal in shaping cellular identity and pathology and has a complex multilevel regulation. Our focus lies on transcription factors (TFs), specifically delving into TF interactions with direct target genes through binding in their promoters or other cis-regulatory regions. Existing methods for building direct target regulons typically lack cell-type specificity or indirectly factor in cell-specific transcript expression. Furthermore, many of the more sophisticated approaches are complicated to implement and apply to new data sets and cell types of interest. To tackle this, we introduce ChIP-Seq-based approaches that, unlike current methods, provide straightforward regulon annotation that includes an additional step to consider isoform-level gene expression.

In this study, we constructed regulons encompassing hundreds of transcription factors (TFs) within four frequently utilized cell lines. Our analysis revealed the enrichment of our regulons in well-established biological networks, and in transcription factor binding motifs. To address the challenge of potential false positive interactions, we explored various filtering strategies and discerned that the open chromatin status proved advantageous for shorter-distance methods (S2Kb and M2Kb).

The comparison of the proposed approaches with existing databases demonstrated a reasonable level of agreement, generally yielding comparable performance to similar data-driven regulons in predicting the effects of TF knockout. While our methods did not surpass state-of-the-art text-mining-based methods (6), the overall performance was similar. A distinguishing advantage of our approach lies in its simplicity and practical utility, relying solely on RNA-Seq and ChIP-Seq data. What differentiates our approach from other ChIP-Seq-based methods is the additional consideration of the transcriptional profile of potential target genes, which enhanced performance in specific cases (such as K-562) during benchmarking.

To exemplify the value of our regulons for context-specific exploration, we conducted case studies involving three cancer scRNA-Seq datasets. Our results illustrated the capacity of our regulons to identify well-known cancer-promoting regulatory programs, highlighting their ability to capture meaningful insights from complex biological data.

However, it is important to acknowledge that the constructed regulons are likely to contain a significant number of false positive interactions, which can arise from various factors including the inherent noise in ChIP-Seq data (20, 56, 57) and nonfunctional TF binding that does not impact transcription. Potential false negatives stem from indirect binding and TF cooperation. Another notable consideration when relying on ChIP-Seq data is the limited coverage across cell lines, with additional biases toward well-studied and prominent TFs. Finally, a key constraint of our proposed methods is that they link each potential TF binding site to the nearest gene, primarily in close proximity. For the S2Kb and M2Kb approaches this limitation leads to the omission of longer-range interactions involving cis-regulatory elements farther away from the target gene. Although the S2Mb approach can incorporate these interactions, they are likely to be false positive associations due to the possibility of a TSS of another gene being closer to the peak than the actual target. A potential future extension would be to use more sophisticated data or predictions of enhancer target genes, for instance using the ABC model (58).

Another important aspect is the consistency of regulons across approaches. Previous research indicates that regulons obtained through diverse methods exhibit limited overlap in identified interactions (5). Here, we similarly observed limited consistency between regulons derived from different methods. This might be attributed to several factors: (1) differences in interaction data sources, where some regulons utilise a single source while others incorporate multiple data sources and modalities, (2) variations in cellular resolution between generalized and cell-type specific regulons, and (3) exclusion or inclusion of indirect TF targets (Table 5).

**Table 5:** Resources of TF-gene interactions utilized for comparison.

| Database | N TFs | Number of interactions in regulons | Annotation process | CT** | Data source | Raw data sources |
| --- | --- | --- | --- | --- | --- | --- |
| GTRD* | 852 (regulons provided for 502 TFs) | 256,331 | TFBSs in the region [-1000,+500] nt around the TSS | - | ChIP-Seq, ChIP-exo, ChIP-nexus, MNase-seq, DNase-seq, FAIRE-seq, ATAC-Seq, RNA-Seq | GEO, SRA, ENCODE, modENCODE |
| ChIP-Atlas | 1807 | K-562: 1,529,999<br>Hep-G2: 2,068,087<br>MCF-7: 639,027<br>GM-12878: 655,040 | Peaks located around ($\pm 1, 5, \text{ or } 10 \text{ kb}$) transcription start sites (TSSs) of RefSeq coding genes | + | ChIP-Seq, ATAC-Seq, DNase-seq, Bisulfite-seq | NCBI SRA |
| DoRothEA | 1541 | 1,076,628 | Literature-curated resources; closest gene to TFBS from ChIP-Seq; TFBS predictions in the promoter; co-expression networks | - | Literature, ChIP-Seq, TFBS motifs, RNA-Seq | Diverse; see (14) |
| CollecTri | 1183 | 45,856 | Multi-source text-mining-based dataset | - | Article abstracts | MedLine, GOA, IntAct, TRRUST, CytReg, GEREDB and SIGNOR |
| TRRUST | 800 | 8,444 | Abstract-based text mining followed by manual curation | - | Article abstracts | MedLine |
| RegNetwork | 1456 | 369,277 | Interactions documented in TRED and KEGG; TFBS predictions in the promoters; PPI pairs that contain at least one TF; miRNA | - | Literature, TFBS motifs, PPIs, miRNA | TRED, KEGG, JASPAR, TRANSFAC, BioGrid, IntAct, KEGG, STRING, HPRD, miRNA databases |

\*Database is cell-type specific, but target gene sets submitted to MSigDB are not
\*\*CT refers to cell-type specificity of a regulon

During this study, we employed a systematic benchmarking process to assess the quality and predictive power of the constructed regulons (19, 40). This approach greatly facilitated comparability across different databases, yet it remains limited due to the relatively small number of covered transcription factors and cell lines (6). For example, in Hep-G2 cells, there were only three knockout experiments available for the 104 shared targets among all tested regulons in the KnockTF database (19). This sparsity of data might contribute to the considerable variation in the predictive capability of a single approach across various cell lines.

Lastly, while our regulons proved valuable for identifying transcription factor activity dysregulation in cancer, a notable drawback is their lack of information regarding the mode and quantitative strength of interactions, such as partial activation or repression. This introduces uncertainty into the sign of TF activity estimates and complicates the interpretation of observed dysregulations. The CollecTri regulons address this by distinguishing between activators and repressors, enhancing reliability for activity estimation. Nevertheless, quantitative assessment of TF regulatory strength remains a challenge. Since motif enrichment-based measures did not yield improved predictive power (6), an intriguing avenue for exploration could involve estimating these measures from the Hills equation based on co-expression networks or direct TF dosage-titration experiments (Domingo et al. in prep; (59)).

In summary, we provided a straightforward approach for annotating TF regulons from cell-type specific TF binding and transcriptional profiles and applied it to four distinct cell lines. We benchmarked these regulons against existing databases using TF knockout experiments and showcased their ability to identify cancer-related dysregulations, highlighting some situations where cell type-specific regulons provided additional information compared to generalized approaches.

## Data Availability

Raw and processed data files are available to download at https://doi.org/10.5281/zenodo.10395639.

Accompanying code is available to review and download at https://github.com/LappalainenLab/chip_seq_regulons

## Acknowledgements

We would like to thank John Morris, and the current and former members of the Lappalainen Lab for helpful discussions and code sharing. Part of the computations were enabled by resources provided by the Swedish National Infrastructure for Computing (SNIC) at UPPMAX partially funded by the Swedish Research Council through grant agreement no. 2018-05973.

