## Supplementary Data for "Specifying cellular context of transcription factor regulons for exploring context-specific gene regulation programs"

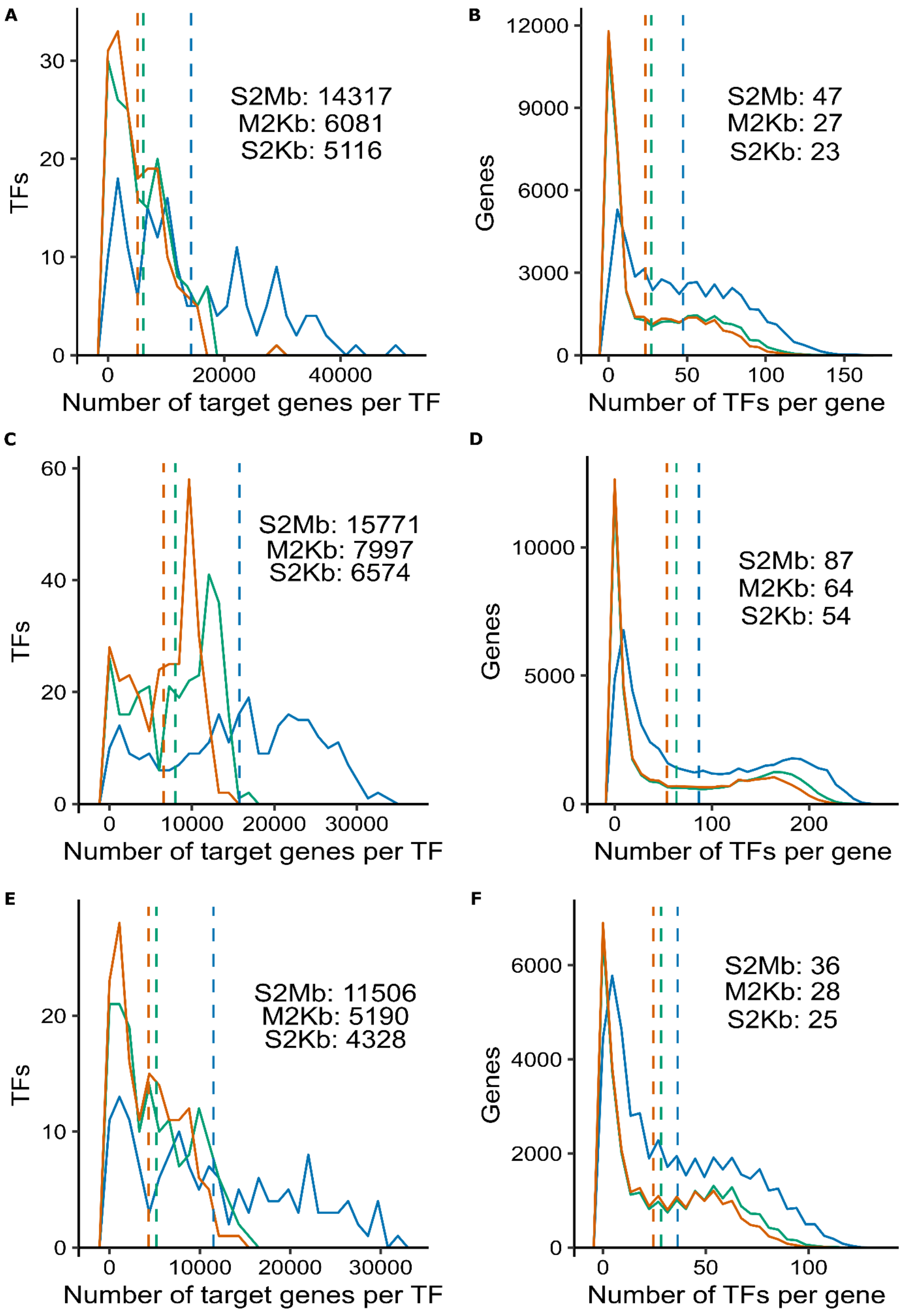

**Figure S1**: Characteristics of regulons, similar to Fig. 1B-C: (A), (C), (E) Distributions of a number of target genes per TF; dashed line shows per method median. (B), (D), (F) Distribution of a number of TFs per target gene; dashed line shows per method median for (A) - (B) MCF-7 regulon, (C) - (D) Hep-G2 regulon, (E) - (F) GM-12878 regulon.

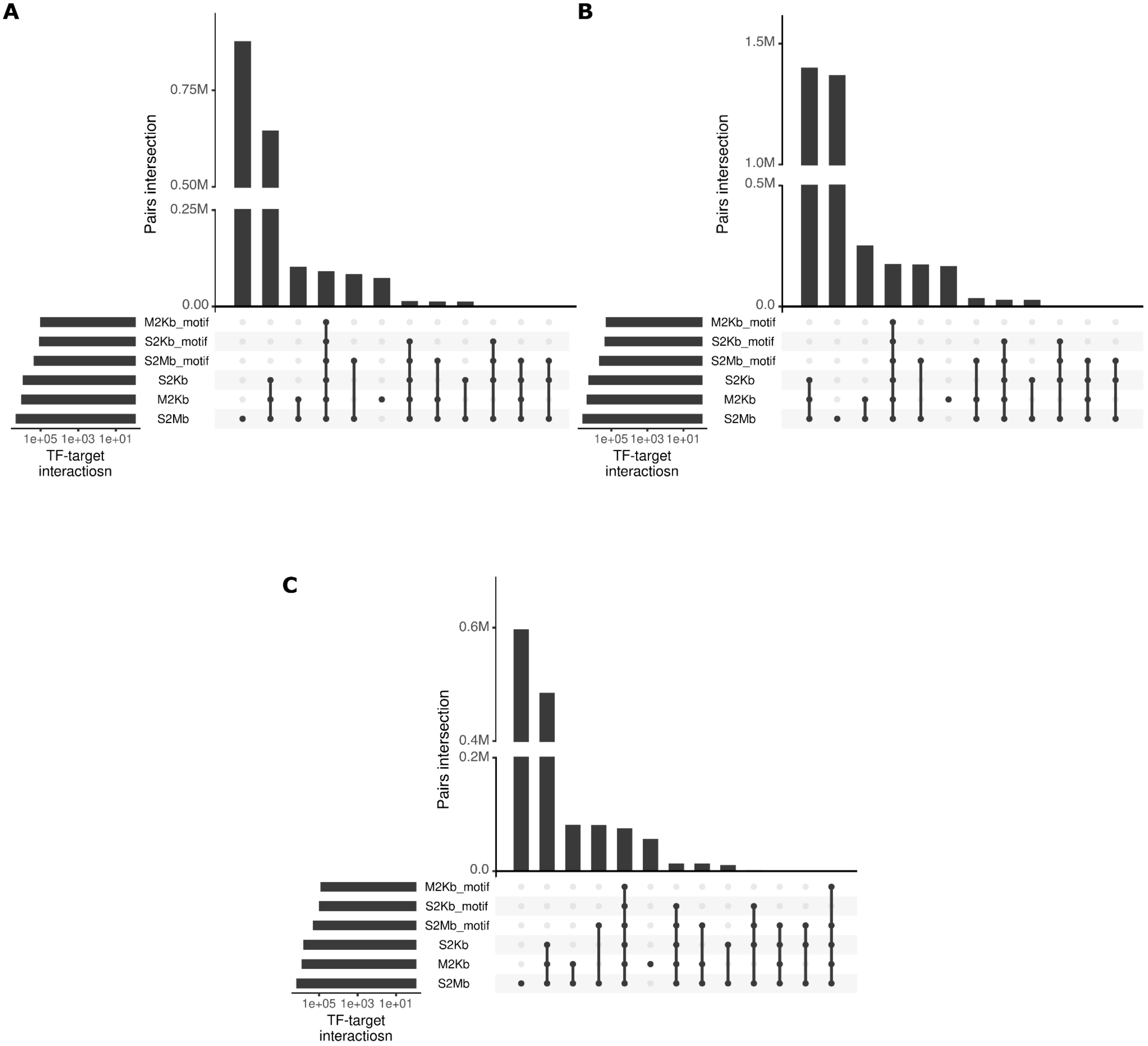


**Figure S2:** Overlap of regulons with TF binding motifs, similar to Fig. 1D. (A) MCF-7 cell line, (B) Hep-G2 cell line, (C) GM-12878 cell line; for ease of interpretation, the top 12 overlapping groups are shown.

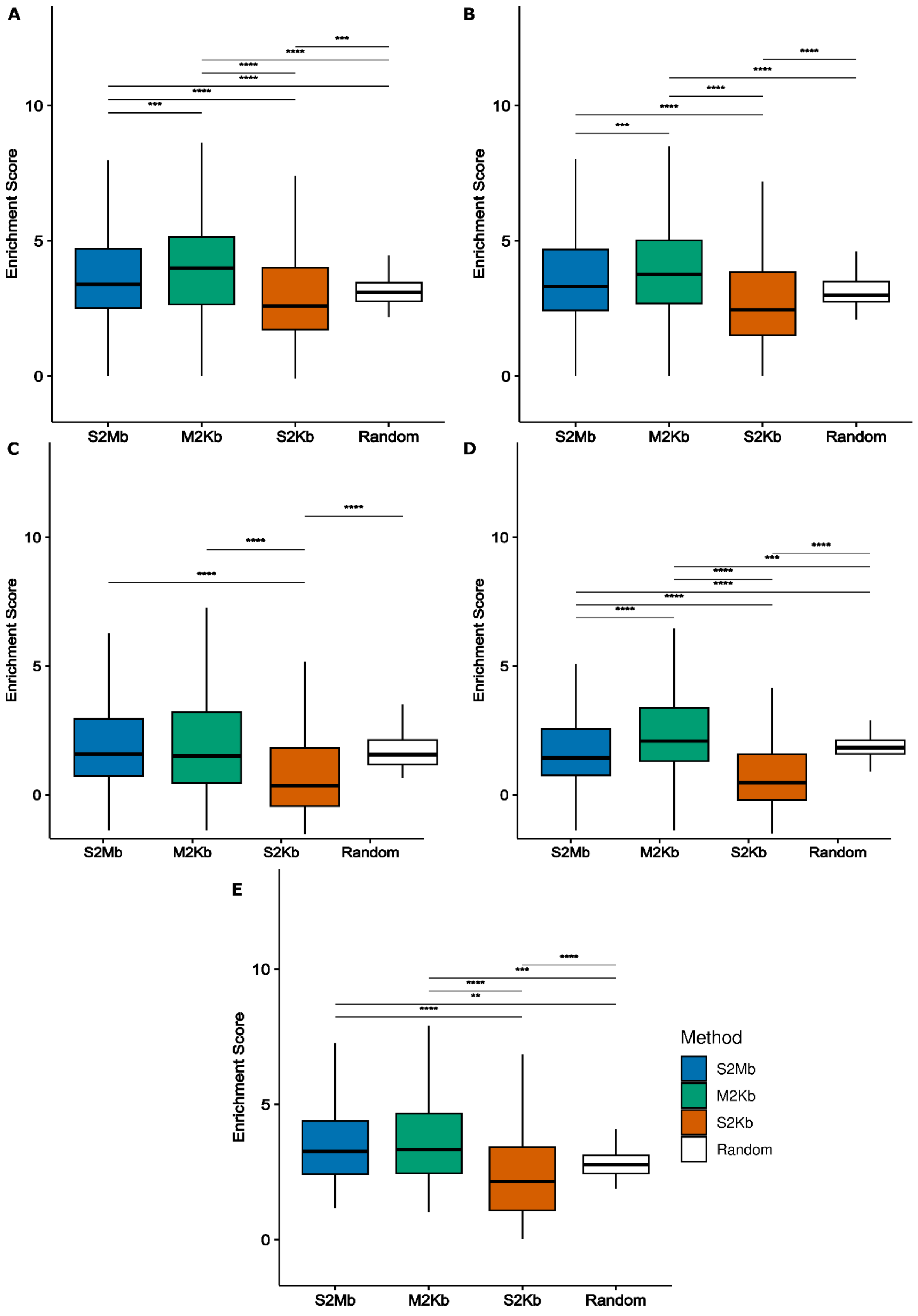
A
B
C
D
E
Enrichment Score
0
5
10
S2Mb
M2Kb
S2Kb
Random
Method
S2Mb
M2Kb
S2Kb
Random

**Figure S3:** Enrichment of regulons in biological networks. (A) Enrichment of interactions from K-562 regulon in coexpression network derived from scRNA-Seq experiment (1). (B) - (E) Enrichment of interactions in protein-protein interaction networks from STRING (2) for (B) K-562, (C) MCF-7, (D) GM-12878 and (E) Hep-G2 regulons. Wilcoxon test with FDR correction is used to determine statistical significance.

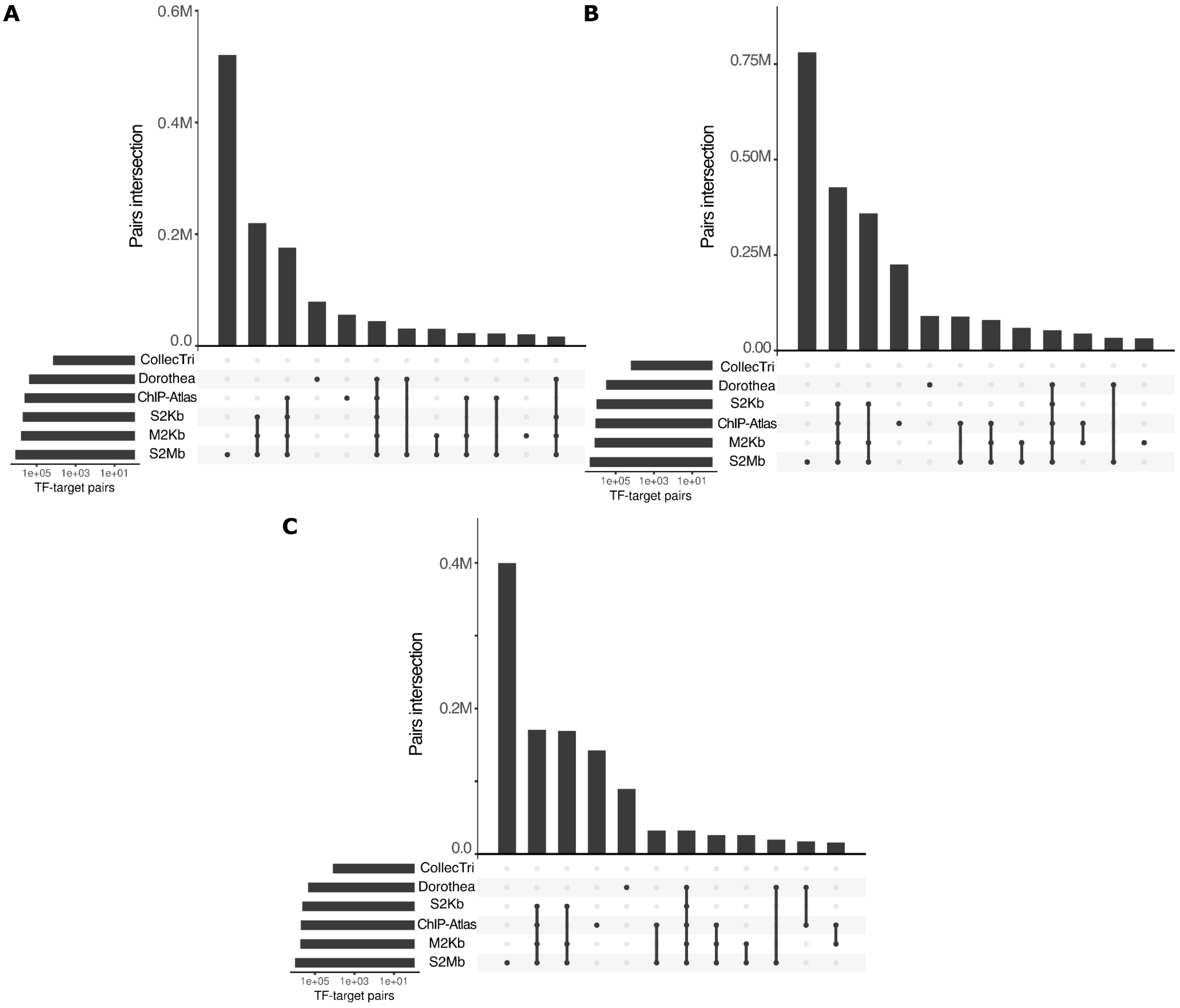


**Figure S4:** Comparison of different regulons, similar to Fig. 2D. Overlap in the TF-target gene pairs between regulons from different resources for (A) MCF-7, (B) Hep-G2 and (C) GM-12878 cell lines; for ease of interpretation, the top 12 overlapping groups are shown.

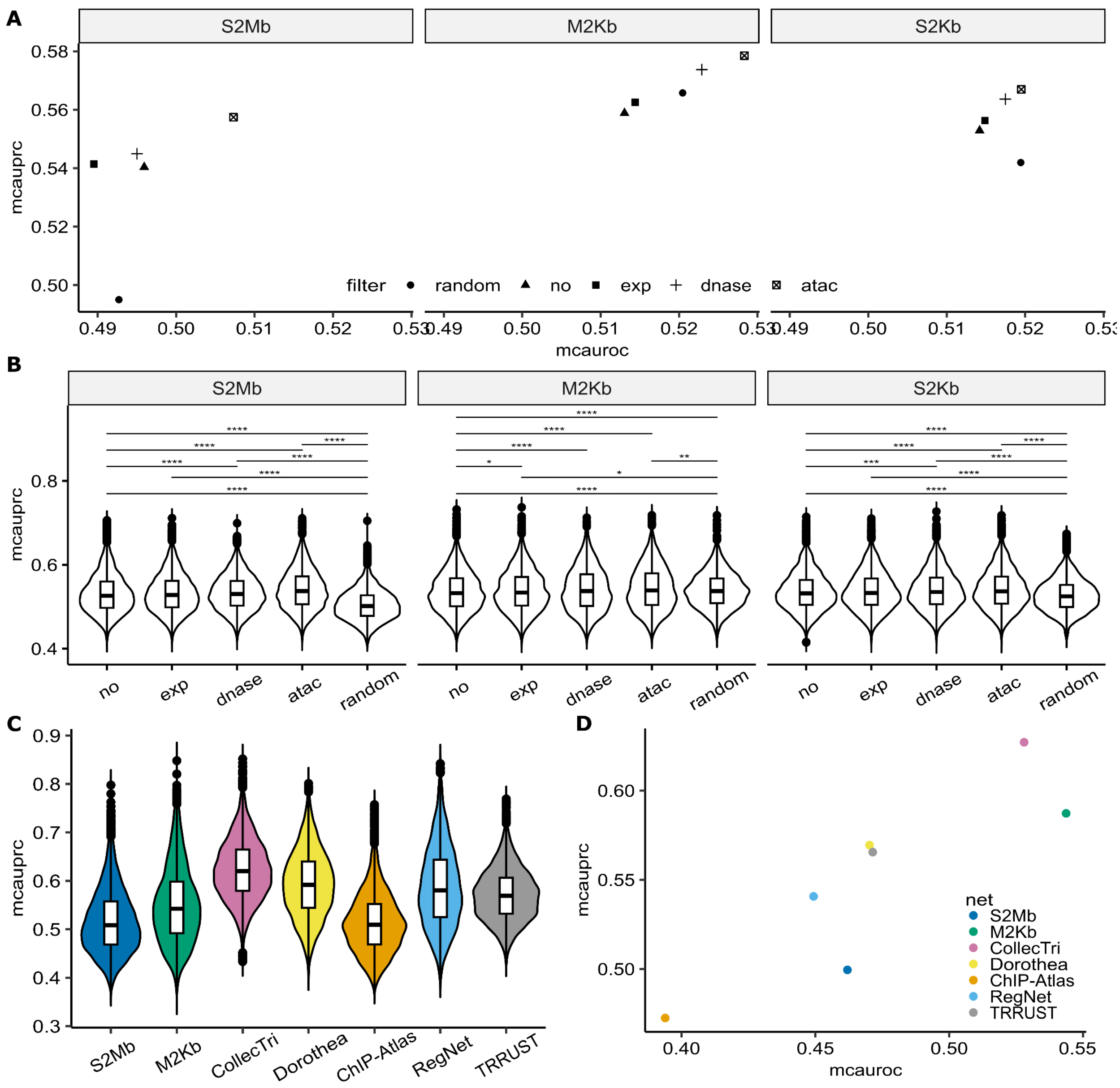


**Figure S5:** Benchmarking of K-562 regulon using knockout experiments from KnockTF database: (A) Impact of filtering strategy on regulon's performance in predicting TF knockout measured by mean MCAUROC (y-axis) and mean MCAUPRC (x-axis) for S2Mb (right), M2Kb (middle), and S2Kb (left) approaches. (B) Distribution of MCAUPRC for filtered regulons for S2Mb (right), M2Kb (middle), and S2Kb (left) approaches. (C) Comparison of the predictive power of regulons using MCAUPRC distribution. (D) Performance regulons in predicting TF knockout measured by mean MCAUROC (y-axis) and mean MCAUPRC (x-axis).

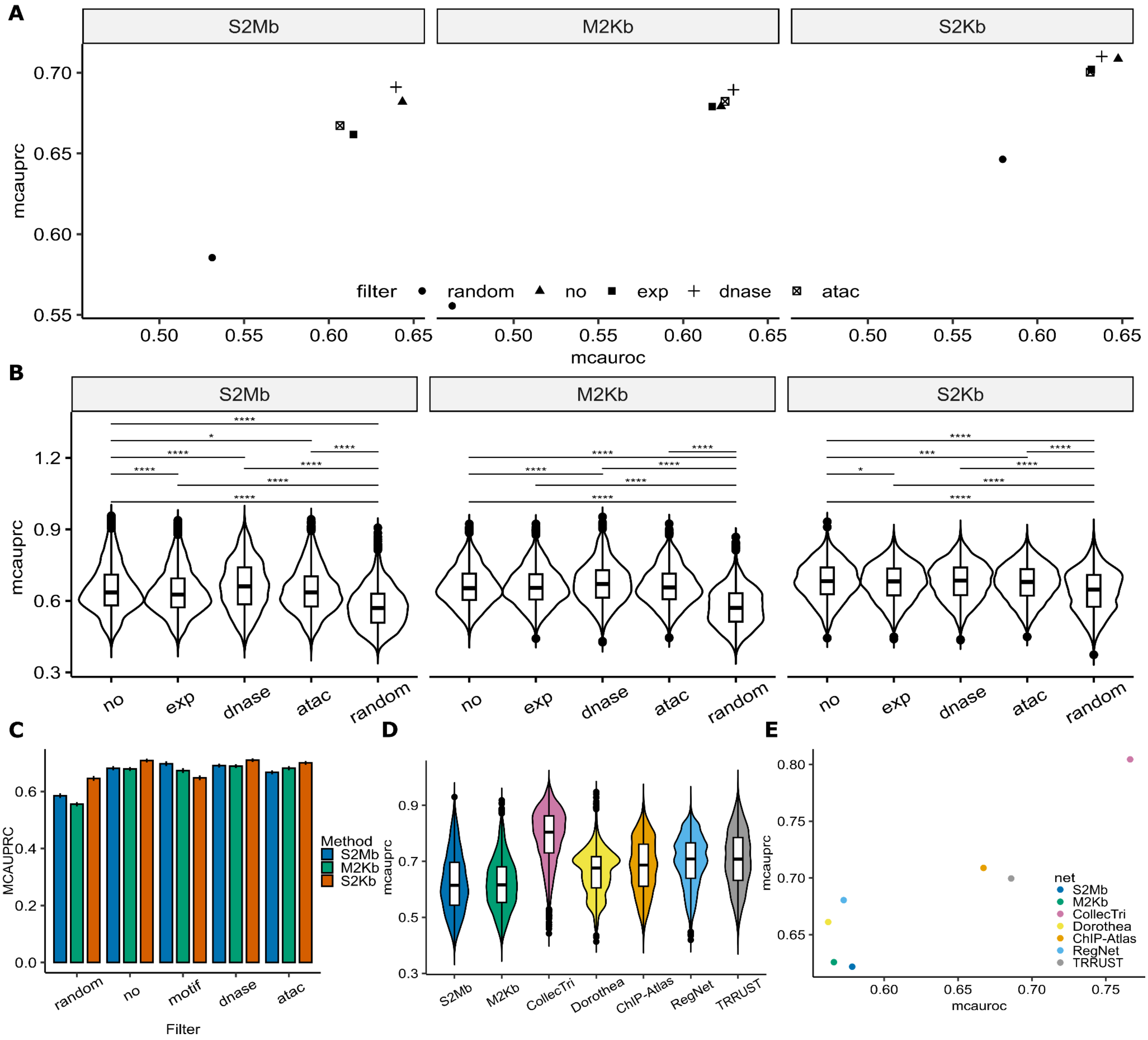


**Figure S6:** Benchmarking of MCF-7 regulon using knockout experiments from KnockTF database, similar to Fig. S5: (A) Impact of filtering strategy on regulon's performance in predicting TF knockout measured by mean MCAUROC (y-axis) and mean MCAUPRC (x-axis) for S2Mb (right), M2Kb (middle), and S2Kb (left) approaches. (B) Distribution of MCAUPRC for filtered regulons for S2Mb (right), M2Kb (middle), and S2Kb (left) approaches. (C) Comparison of filtering strategies for S2Mb (in blue), M2Kb (in red), and S2Kb (in green). (D) Comparison of the predictive power of regulons using MCAUPRC distribution. (E) Performance regulons in predicting TF knockout measured by mean MCAUROC (y-axis) and mean MCAUPRC (x-axis).

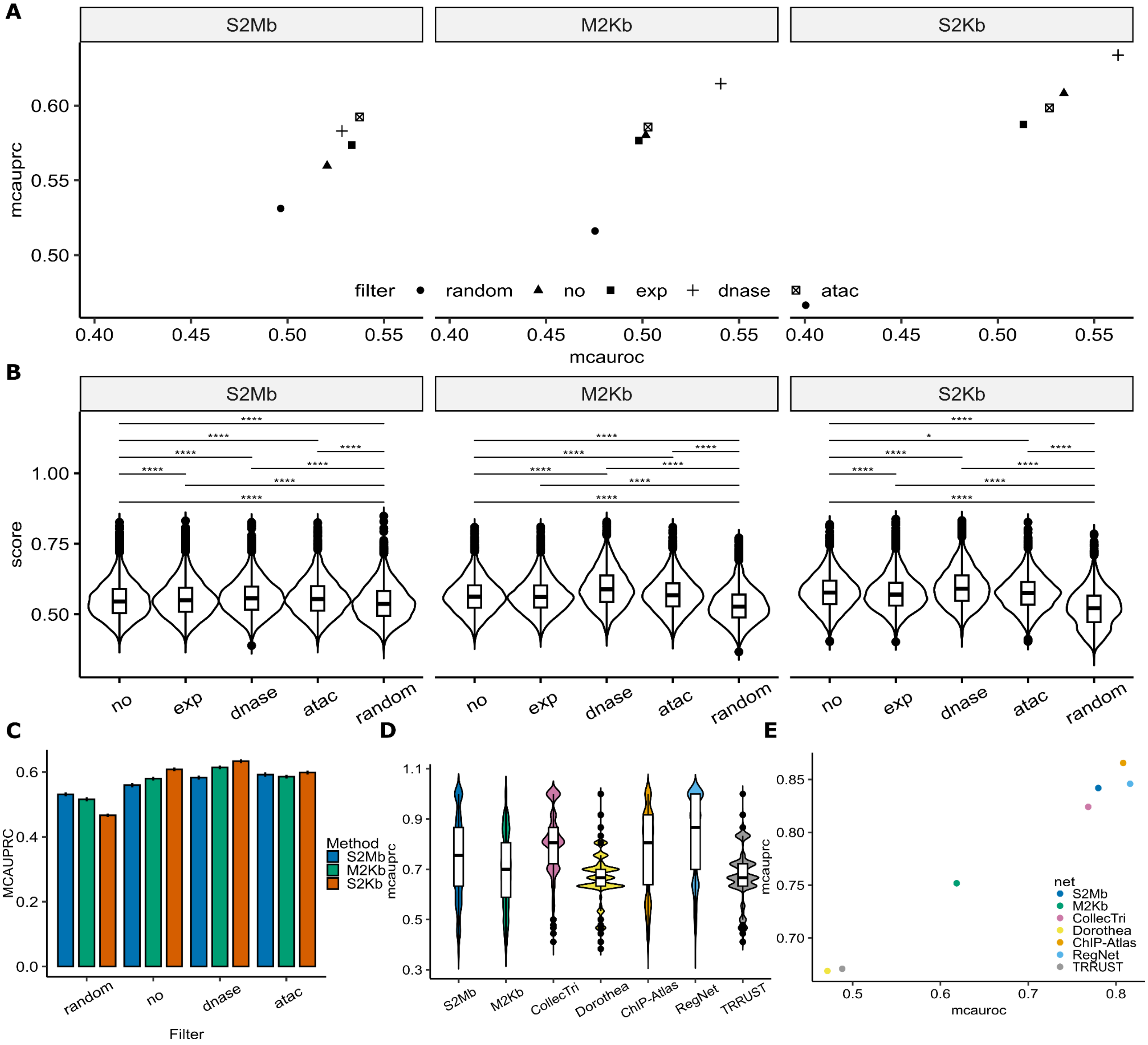


**Figure S7:** Benchmarking of Hep-G2 regulon using knockout experiments from KnockTF database, similar to Fig. S5: (A) Impact of filtering strategy on regulon's performance in predicting TF knockout measured by mean MCAUROC (y-axis) and mean MCAUPRC (x-axis) for S2Mb (right), M2Kb (middle), and S2Kb (left) approaches. (B) Distribution of MCAUPRC for filtered regulons for S2Mb (right), M2Kb (middle), and S2Kb (left) approaches. (C) Comparison of filtering strategies for S2Mb (in blue), M2Kb (in red), and S2Kb (in green). (D) Comparison of the predictive power of regulons using MCAUPRC distribution. (E) Performance regulons in predicting TF knockout measured by mean MCAUROC (y-axis) and mean MCAUPRC (x-axis).

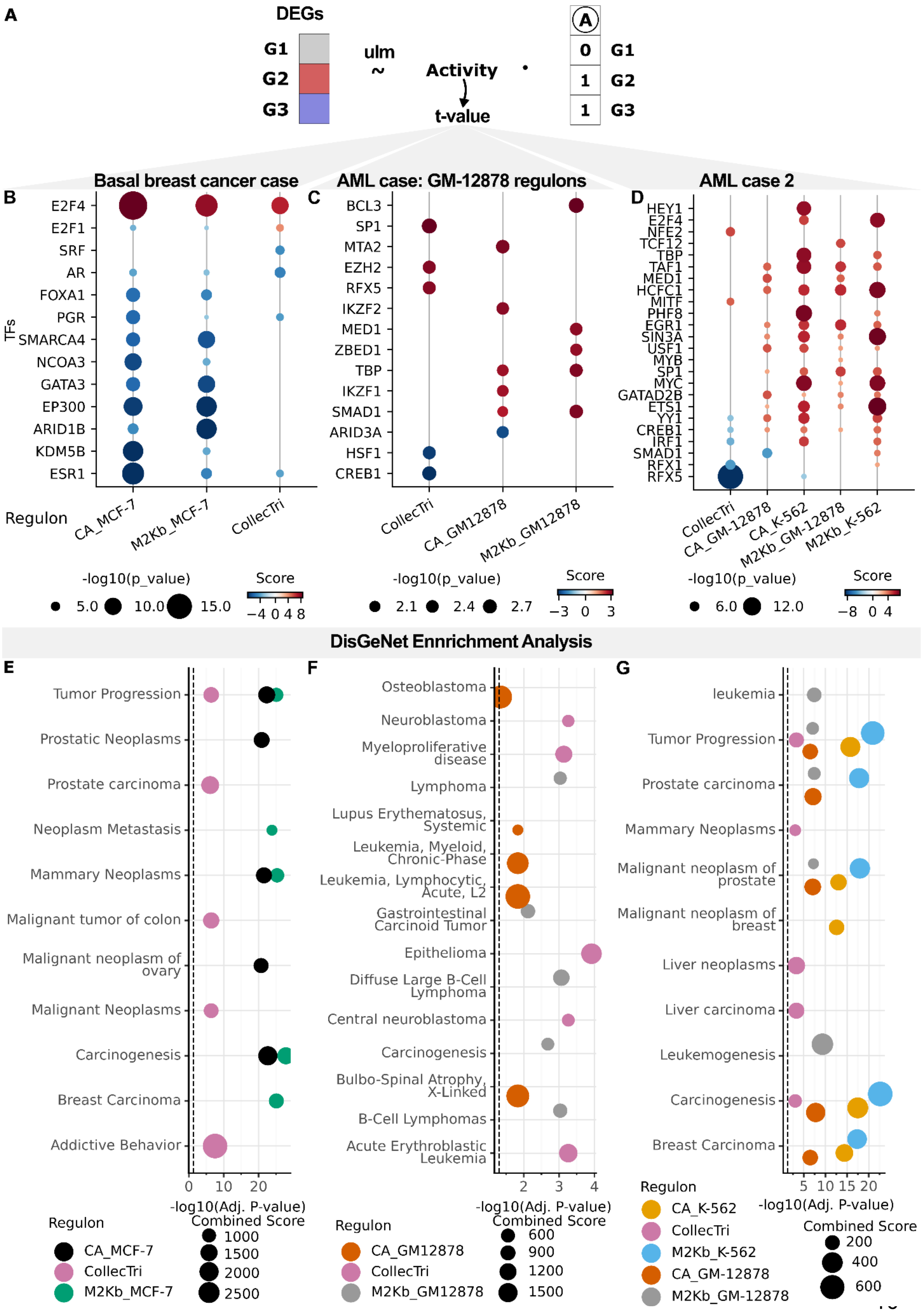
A
DEGs
G1
G2
G3
ulm
~
Activity
t-value
A
0
1
1
G1
G2
G3
Basal breast cancer case
AML case: GM-12878 regulons
AML case 2
B
E2F4
E2F1
SRF
AR
FOXA1
PGR
SMARCA4
NCOA3
GATA3
EP300
ARID1B
KDM5B
ESR1
TFs
Regulon
CA_MCF-7
M2Kb_MCF-7
CollecTri
C
BCL3
SP1
MTA2
EZH2
RFX5
IKZF2
MED1
ZBED1
TBP
IKZF1
SMAD1
ARID3A
HSF1
CREB1
CollecTri
CA_GM12878
M2Kb_GM12878
D
HEY1
E2F4
NFE2
TCF12
TBP
TAF1
MED1
HCFC1
MITF
PHF8
EGR1
SIN3A
USF1
MYB
SP1
MYC
GATAD2B
ETS1
YY1
CREB1
IRF1
SMAD1
RFX1
RFX5
CollecTri
CA_GM-12878
CA_K-562
M2Kb_GM-12878
M2Kb_K-562
-log10(p_value)
5.0
10.0
15.0
Score
−4 0 4 8
-log10(p_value)
2.1
2.4
2.7
Score
−3 0 3
-log10(p_value)
6.0
12.0
Score
−8 0 4
DisGeNet Ennrichment Analysis
E
Tumor Progression
Prostatic Neoplasms
Prostate carcinoma
Neoplasm Metastasis
Mammary Neoplasms
Malignant tumor of colon
Malignant neoplasm of ovary
Malignant Neoplasms
Carcinogenesis
Breast Carcinoma
Addictive Behavior
0 10 20
-log10(Adj. P-value)
Regulon
CA_MCF-7
CollecTri
M2Kb_MCF-7
Combined Score
1000
1500
2000
2500
F
Osteoblastoma
Neuroblastoma
Myeloproliferative disease
Lymphoma
Lupus Erythematosus, Systemic
Leukemia, Myeloid, Chronic-Phase
Leukemia, Lymphocytic, Acute, L2
Gastrointestinal Carcinoid Tumor
Epithelioma
Diffuse Large B-Cell Lymphoma
Central neuroblastoma
Carcinogenesis
Bulbo-Spinal Atrophy, X-Linked
B-Cell Lymphomas
Acute Erythroblastic Leukemia
2 3 4
-log10(Adj. P-value)
Regulon
CA_GM12878
CollecTri
M2Kb_GM12878
Combined Score
600
900
1200
1500
G
leukemia
Tumor Progression
Prostate carcinoma
Mammary Neoplasms
Malignant neoplasm of prostate
Malignant neoplasm of breast
Liver neoplasms
Liver carcinoma
Leukemogenesis
Carcinogenesis
Breast Carcinoma
5 10 15 20
-log10(Adj. P-value)
Regulon
CA_K-562
CollecTri
M2Kb_K-562
CA_GM-12878
M2Kb_GM-12878
Combined Score
200
400
600

**Figure S8:** Case studies of detecting transcriptional dysregulation in cancers, similar to Fig. 3. (A) Schematic overview of activity estimation procedure. (B) - (D) Activity estimate for the top-ranked and manually selected TFs of each regulon for (B) basal type breast cancer (3), (C) leukemic progenitors (4) using GM-12878 regulons and (D) hematopoietic stem cells (4). Here positive and negative activity scores mean activation and deactivation of TFs in malignant cells respectively. (E) - (G) Top 5 enriched terms of the enrichment analysis of dysregulated TFs in disease gene from DisGeNet database (5) for (E) basal type breast cancer, (F) leukemic progenitors and (G) hematopoietic stem cells. Here "CA_cell line" refers to ChIP-Atlas regulons for the specified cell line and "M2Kb_cell line" refers to M2Kb regulons.

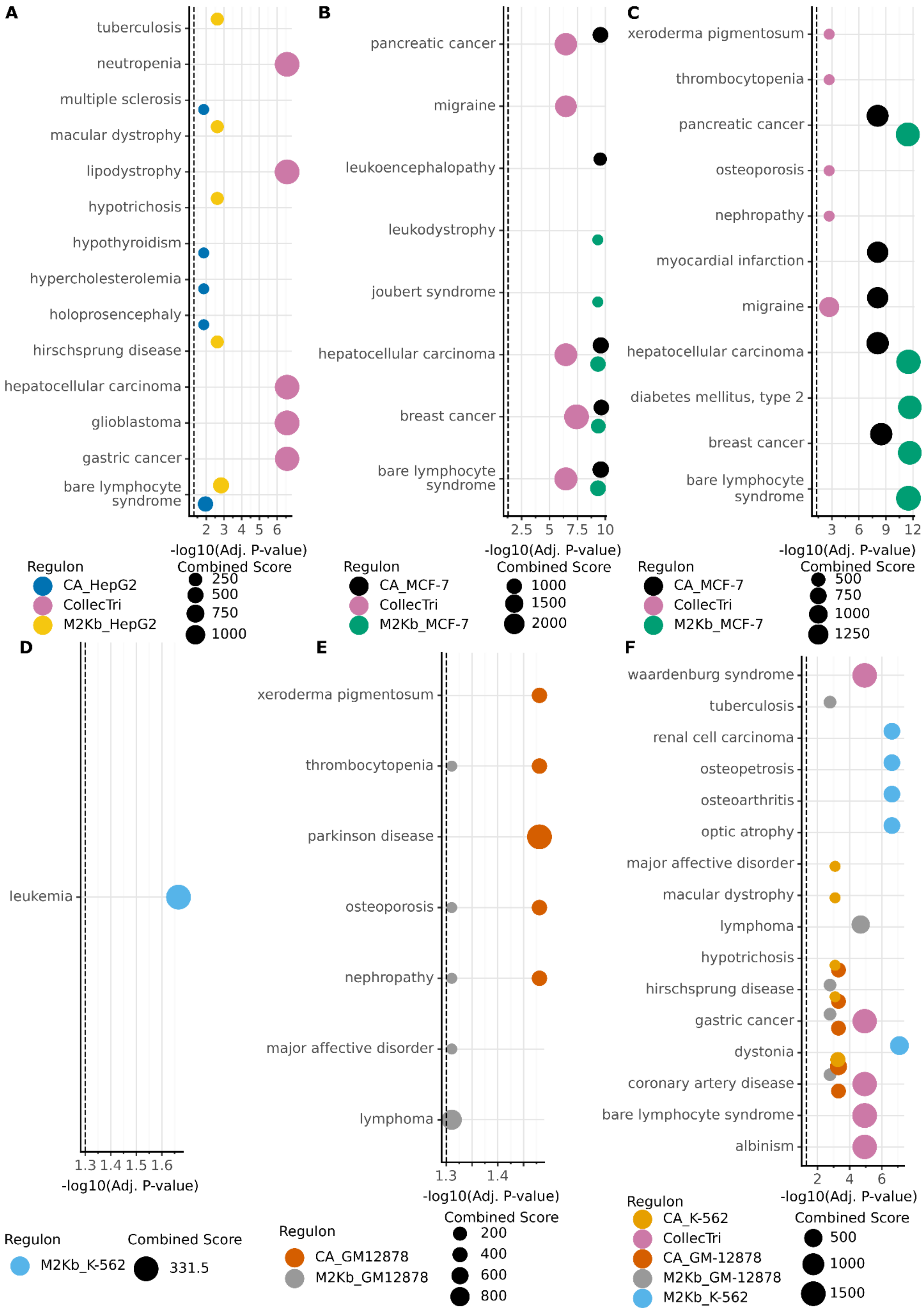
A
tuberculosis
neutropenia
multiple sclerosis
macular dystrophy
lipodystrophy
hypotrichosis
hypothyroidism
hypercholesterolemia
holoprosencephaly
hirschsprung disease
hepatocellular carcinoma
glioblastoma
gastric cancer
bare lymphocyte syndrome
2 3 4 5 6
-log10(Adj. P-value)
Regulon
CA_HepG2
CollecTri
M2Kb_HepG2
Combined Score
250
500
750
1000
B
pancreatic cancer
migraine
leukoencephalopathy
leukodystrophy
joubert syndrome
hepatocellular carcinoma
breast cancer
bare lymphocyte syndrome
2.5 5 7.5 10
-log10(Adj. P-value)
Regulon
CA_MCF-7
CollecTri
M2Kb_MCF-7
Combined Score
1000
1500
2000
C
xeroderma pigmentosum
thrombocytopenia
pancreatic cancer
osteoporosis
nephropathy
myocardial infarction
migraine
hepatocellular carcinoma
diabetes mellitus, type 2
breast cancer
bare lymphocyte syndrome
3 6 9 12
-log10(Adj. P-value)
Regulon
CA_MCF-7
CollecTri
M2Kb_MCF-7
Combined Score
500
750
1000
1250
D
leukemia
1.3 1.4 1.5 1.6
-log10(Adj. P-value)
Regulon
M2Kb_K-562
Combined Score
331.5
E
xeroderma pigmentosum
thrombocytopenia
parkinson disease
osteoporosis
nephropathy
major affective disorder
lymphoma
1.3 1.4
-log10(Adj. P-value)
Regulon
CA_GM12878
M2Kb_GM12878
Combined Score
200
400
600
800
F
waardenburg syndrome
tuberculosis
renal cell carcinoma
osteopetrosis
osteoarthritis
optic atrophy
major affective disorder
macular dystrophy
lymphoma
hypotrichosis
hirschsprung disease
gastric cancer
dystonia
coronary artery disease
bare lymphocyte syndrome
albinism
2 4 6
-log10(Adj. P-value)
Regulon
CA_K-562
CollecTri
CA_GM-12878
M2Kb_GM-12878
M2Kb_K-562
Combined Score
500
1000
1500

**Figure S9:** Enrichment analysis of dysregulated TFs in disease gene from OMIM database (6). (A) hepatoblastoma (7), (B) lumenal type A breast cancer (3), (C) basal type breast cancer (3), leukemic progenitors using K-562 (D) or GM-12878 regulons (E) and (F) hematopoietic stem cells (4). Here “CA_cell line” refers to ChIP-Atlas regulons for the specified cell line and “cell line” refers to M2Kb regulons.

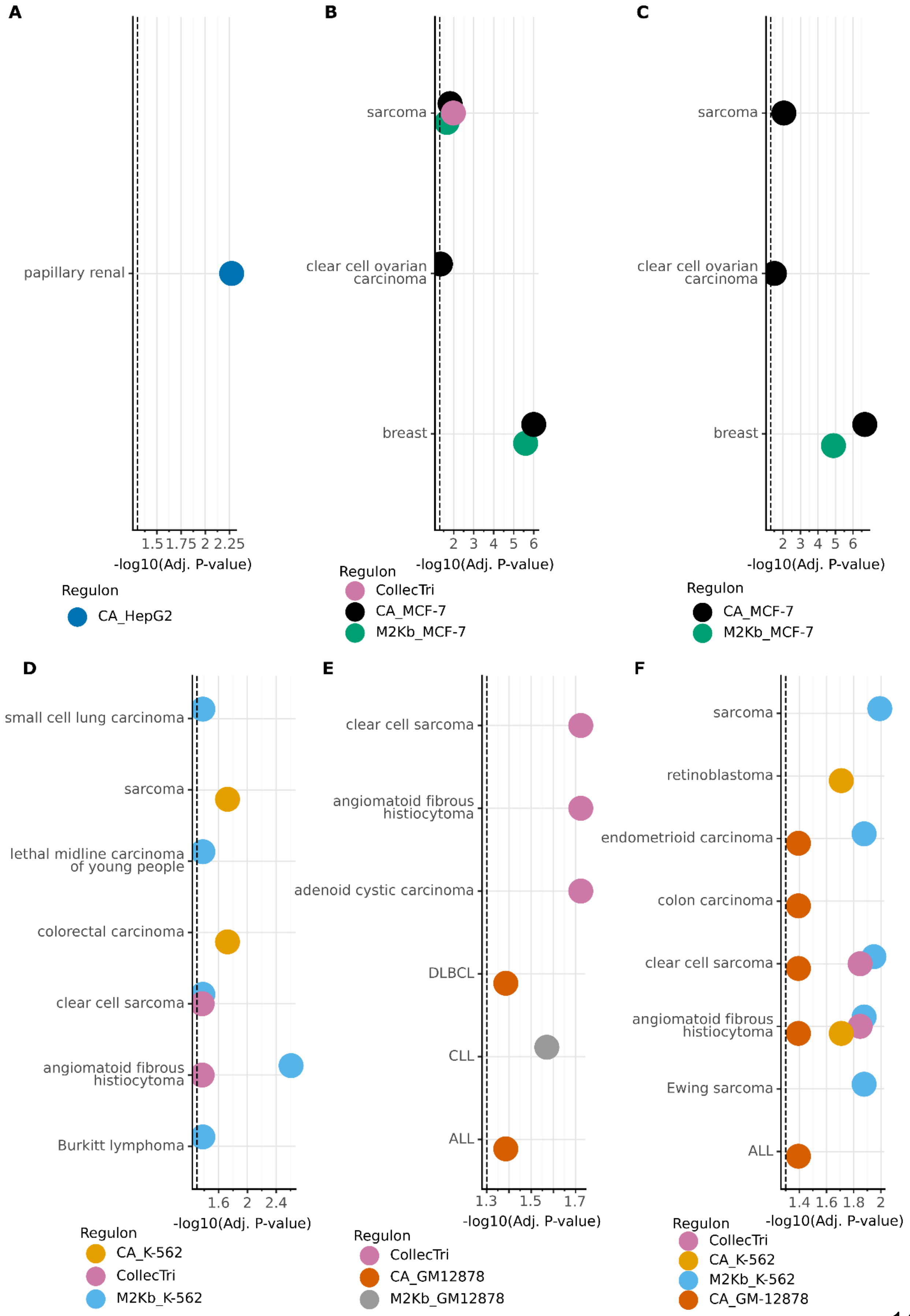
A
papillary renal
1.5 1.75 2 2.25
-log10(Adj. P-value)
Regulon
CA_HepG2
B
sarcoma
clear cell ovarian carcinoma
breast
2 3 4 5 6
-log10(Adj. P-value)
Regulon
CollecTri
CA_MCF-7
M2Kb_MCF-7
C
sarcoma
clear cell ovarian carcinoma
breast
2 3 4 5 6
-log10(Adj. P-value)
Regulon
CA_MCF-7
M2Kb_MCF-7
D
small cell lung carcinoma
sarcoma
lethal midline carcinoma of young people
colorectal carcinoma
clear cell sarcoma
angiomatoid fibrous histiocytoma
Burkitt lymphoma
1.6 2 2.4
-log10(Adj. P-value)
Regulon
CA_K-562
CollecTri
M2Kb_K-562
E
clear cell sarcoma
angiomatoid fibrous histiocytoma
adenoid cystic carcinoma
DLBCL
CLL
ALL
1.3 1.5 1.7
-log10(Adj. P-value)
Regulon
CollecTri
CA_GM12878
M2Kb_GM12878
F
sarcoma
retinoblastoma
endometrioid carcinoma
colon carcinoma
clear cell sarcoma
angiomatoid fibrous histiocytoma
Ewing sarcoma
ALL
1.4 1.6 1.8 2
-log10(Adj. P-value)
Regulon
CollecTri
CA_K-562
M2Kb_K-562
CA_GM-12878

**Figure S10:** Enrichment analysis of dysregulated TFs in consensus gene from COSMIC database (8), similar to Fig. S9. (A) hepatoblastoma (7), (B) lumenal type A breast cancer (3), (C) basal type breast cancer (3), leukemic progenitors using K-562 (D) or GM-12878 regulons (E) and (F) hematopoietic stem cells (4). Here “CA_cell line” refers to ChIP-Atlas regulons for the specified cell line and “cell line” refers to M2Kb regulons.

**Supplementary Table 1:** Top 5 KEGG pathways enriched in dysregulated TFs in liver cancer detected using M2Kb_HepG2 regulon

| Term | Overlap | Adjusted P-value | Odds Ratio | Combined Score |
|---|---|---|---|---|
| Transcriptional misregulation in cancer | 15/192 | 3.86E-11 | 16.87 | 487.4 |
| Human T-cell leukemia virus 1 infection | 12/219 | 3.09E-07 | 11.18 | 214.8 |
| Hepatitis B | 8/162 | 1.88E-04 | 9.67 | 119.9 |
| Antigen processing and presentation | 6/78 | 1.88E-04 | 15.29 | 185.1 |
| Cellular senescence | 7/156 | 7.31E-04 | 8.67 | 89.8 |

**Supplementary Table 2:** Top 5 KEGG pathways enriched in dysregulated TFs in liver cancer detected using CollecTri regulon

| Term | Overlap | Adjusted P-value | Odds Ratio | Combined Score |
|---|---|---|---|---|
| Pathways in cancer | 25/531 | 3.14E-20 | 22.32 | 1114.2 |
| Human T-cell leukemia virus 1 infection | 17/219 | 5.14E-17 | 32.56 | 1361.7 |
| Th17 cell differentiation | 13/107 | 1.70E-15 | 49.88 | 1891.5 |
| Hepatitis B | 13/162 | 3.17E-13 | 31.38 | 1016.9 |
| Osteoclast differentiation | 12/127 | 4.09E-13 | 36.93 | 1178.9 |
